# Micro-state-based neural decoding of speech categorization using Bayesian non-parametrics

**DOI:** 10.1101/2021.11.17.469011

**Authors:** Rakib Al-Fahad, Mohammed Yeasin, Kazi Ashraf Moinuddin, Gavin M Bidelman

## Abstract

Understanding the many-to-many mapping between patterns of functional brain connectivity and discrete behavioral responses is critical for speech-language processing. We present a microstate-based analysis of EEG recordings to characterize spatio-temporal dynamics of neural activities that underly rapid speech categorization decisions. We implemented a data driven approach using Bayesian non-parametrics to capture the mapping between EEG and the speed of listeners’ phoneme identification [i.e., response time (RT)] during speech labeling tasks. Based on our empirical analyses, we show task-relevant events such as resting-state, stimulus coding, auditory-perceptual object (category) formation, and response selection can be explained using patterns of micro-state dwell-time and are decodable as unique time segments during speech perception. State-dependent activities localize to a fronto-temporo-parietal circuit (superior temporal, supramarginal, inferior frontal gyri) exposing a core decision brain network (DN) underlying rapid speech categorization. Furthermore, RTs were inversely proportional to the frequency of state transitions, such that the rate of change between brain microstates was higher for trials with slower compared to faster RTs. Our findings imply that during rapid speech perception, higher uncertainty producing prolonged RTs (slower decision-making) is associated with staying in the DN longer compared lower RTs (faster decisions). We also show that listeners’ perceptual RTs are highly sensitive to individual differences. Our computational method opens a new avenue in segmentation and dynamic brain connectivity for modeling neuroimaging data and understanding task-related cognitive events.

## INTRODUCTION

The acoustic nature of speech is continuous; even identical utterances can be produced with stark differences in their physical acoustic dimensions (e.g., talker variability in pitch or timbre (Prather, Nowicki, Anderson, Peters, & Mooney, 2009). Yet, speech perception unfolds as a discrete process, invoking a “down-sampling” mechanism that enables listeners to group sounds into smaller sets of (phonetic) categories. This binning process is known as categorical perception (CP) (Harnad & Bureau, 1987; Liberman, Cooper, Shankweiler, & Studdert-Kennedy, 1967; Pisoni, 1973; Pisoni & Luce, 1987) and is fundamental not only to maintaining perceptual constancy of the soundscape but also everyday speech listening.

When identifying speech, listeners’ response times (RTs) provide a window into the speed of their decision process and reveal stark individual differences in perceptual labeling speeds (Bidelman, Moreno, & Alain, 2013; Pisoni & Tash, 1974). For example, listeners categorize prototypical speech sounds (e.g., exemplars from their native language) much faster than ambiguous or less familiar ones (e.g., nonnative speech sounds) (Bidelman & Lee, 2015). RTs also slow near perceptual boundaries, where listeners shift from hearing one linguistic class to another (e.g., /u/ vs. /a/ vowel) and presumably require more time to access the “correct” speech template (Bidelman et al., 2013; Liebenthal et al., 2010; Pisoni & Tash, 1974; Reetzke, Xie, Llanos, & Chandrasekaran, 2018). Rapid speech identification is also highly sensitive to stimulus familiarity (Bidelman & Walker, 2017; Liebenthal et al., 2010; Lively, Logan, & Pisoni, 1993) an individual’s experience (Bidelman & Lee, 2015; Liberman et al., 1967), and certain disorders that affect receptive speech listening skills (Bidelman, Lowther, Tak, & Alain, 2017; Bidelman, Villafuerte, Moreno, & Alain, 2014; Calcus Axelle, Lorenzi Christian, Collet Gregory, Colin Cécile, & Kolinsky Régine, 2016; Hakvoort Britt et al., 2016). Consequently, understanding the neural mechanisms that drive individual differences in categorization might help elucidate not only how continuous sensory information is mapped to discrete, perceptual representations in the brain (Bidelman et al., 2013; Phillips, 2001; Pisoni & Luce, 1987) but also inform putative targets for speech rehabilitation.

While the neural correlates of CP have been well documented in terms of the regional contributions to behavior, our recent study (Al-Fahad, Yeasin, & Bidelman, 2020) took a different approach, examining neural speech processing and CP from a full-brain (functional connectivity) perspective. Using neural decoding and graph mining techniques applied to single-trial EEG, we showed that unique patterns of functional connectivity among a circuit involving superior temporal gyrus (STG), parietal, motor, and prefrontal regions distinguished the speed of listeners’ speech labeling (i.e., RTs). Slow responders tended to utilize the same functional brain networks excessively (or inappropriately) whereas fast responders utilized the same neural pathways but with more restricted organization. While our findings demonstrate the strength of network-level descriptions of the brain map to different behavioral outcomes, they did not consider the time dynamics of brain activity. In fact, the assumption that functional connectivity is static in time (as in most studies) discards important temporal dynamics of EEG which are potentially behaviorally-relevant. As such, previous work cannot speak to the different and evolving brain states that might subserve various processes over the time course of speech identification tasks (e.g., stimulus coding, acoustic-phonetic conversion, lexical interface, response selection, etc.).

Moving toward a more dynamic view of functional brain connectivity (FC), microstate analysis is a recent advance that allows for a time-varying view of neural coupling via the identification of salient transitory states of brain responses (Calhoun, Miller, Pearlson, & Adalı, 2014; Koenig et al., 1999). Coupling refers to possible time-varying levels of correlated or mutually informed activity. Because this technique simultaneously considers signals recorded from all areas of cortex, it is capable of assessing the function of large-scale functional brain networks (Khanna, Pascual-Leone, Michel, & Farzan, 2015). Such networks often referred as microstate functional connectivity (*μ*FC) networks. The *μ*FC is a widely used tool for studying the temporal dynamics of whole-brain FC patterns (Michel & Koenig, 2018). State-of-the-art methods for investigating dynamic FC in EEG recordings mainly follows two common strategies: (i) a temporal sliding window approach (Hindriks et al., 2016; Hutchison et al., 2013; Karamzadeh, Medvedev, Azari, Gandjbakhche, & Najafizadeh, 2013; O’Neill et al., 2017; Preti, Bolton, & Van De Ville, 2017; Shakil, Lee, & Keilholz, 2016) or (ii) an adaptive segmentation via clustering approach (Allen et al., 2014; Damaraju et al., 2014; Hutchison et al., 2013; Mheich, Hassan, Khalil, Berrou, & Wendling, 2015; Shakil et al., 2016). Both firmly rely on the window size, and the strategy for selecting a reasonable window extent remains unsolved. Microstate-based methods also usually follow K-means or Gaussian mixture modeling (GMM) based clustering (Allen et al., 2014; Damaraju et al., 2014; Hutchison et al., 2013; Mheich et al., 2015; Shakil et al., 2016). Despite the usefulness of these clustering-based methods, they have several drawbacks. Choosing a proper number of clusters can be challenging for dynamic data with no prior knowledge. Hard-clustering algorithms are also sensitive to noise and outliers and suffer poor generalization across studies. In addition, these approaches cannot handle an infinite number of clusters and representative microstates are often changed with new observations.

Hierarchical Dirichlet Process HMM (HDP-HMM) (Beal, Ghahramani, & Rasmussen, 2002; Fox, Sudderth, Jordan, & Willsky, 2011; Teh, Jordan, Beal, & Blei, 2005) provides an elegant Bayesian Nonparametric framework for sequential data segmentation with different numbers of states. State-of-the-art inference algorithms for HMMs and HDP-HMMs cannot efficiently learn from large datasets, often getting trapped at local optima and not exploring segmentations with a varying finite number of states (M. C. Hughes, Stephenson, & Sudderth, 2015). In addition, stochastic optimization methods (Foti, Xu, Laird, & Fox, 2014; Johnson & Willsky, 2014) cannot change the number of states during execution, making them vulnerable to large datasets and convergence issues. To overcome these limitations, Monte Carlo procedures (Chang & Fisher III, 2014; Fox, Hughes, Sudderth, & Jordan, 2014; Wang & Blei, 2012) can use the entire dataset, but require all sequences to fit into memory. Consequently, they have the drawback of being computationally inefficient and non-scalable. To avoid these limitations, some studies (i) convert multivariate EEG data into a representative univariate time series [e.g., use Global Field Power (Michel & Koenig, 2018)] or (ii) use trial- or subject-wise averages to reduce dimensionality of the data (Duc & Lee, 2019).

A promising solution to overcome these computational complexities is “Memoized Variational Inference (moVB)” (M. C. Hughes & Sudderth, 2013). Generalizing this algorithm for Dirichlet Process (DP) mixture (M. Hughes, Kim, & Sudderth, 2015) and HDP topic models (M. Hughes et al., 2015), Michael et al. proposed an inference algorithm for the sticky HDP-HMM that scales to big datasets by processing a few sequences at a time (M. C. Hughes et al., 2015). Hence, this algorithm is scalable, reliable, and is fast to converge. The sticky HDP-HMM with moVB enables one to (i) dynamically segment task-related or resting state multivariate EEG, (ii) discover an appropriate number of micro-states, and (iii) use birth, merge, and delete operations (M. C. Hughes et al., 2015) to avoid an infinite number of clusters in the solution. Furthermore, it allows one to update a pretrained model with a new observation, thus allowing for prediction of future data.

Here, we aimed to characterize the temporal dynamics of the brain underlying a core skill for speech perception (i.e., categorization) by discovering patterned states of spatiotemporal neural activity that describe different microstates underlying this critical perceptual-cognitive process. We adopted sticky HDP-HMM with moVB-based dynamic EEG data segmentation analyses to: (i) identify the different brain microstates which unfold as listeners label sounds, (ii) characterize possible differences in the rate of change and/or number (entropy) of state transitions that might distinguish fast from slow perceivers, and (iii) determine the brain areas which describe the decision network (DN) engaged during CP of speech. Our data-driven approach reveals that the duration which listeners stay in the DN is directly related to their perceptual speed, and thus skill level, in rapid speech identification.

## METHODS

### Participants

N=35 adults (12 males, 23 females) were recruited from the University of Memphis student body and Greater Memphis Area to participate in the experiment. All but one was between the age of 18 and 35 years (M = 24.5, SD = 6.9 years). All exhibited normal hearing sensitivity confirmed via audiometric screening (i.e., ¡ 20 dB HL, octave frequencies 250 - 8000 Hz), were strongly right-handed (77.1± 36.4 laterality index [46]), and had obtained at least a collegiate level of education (17.2 ± 2.9 years). None had any history of neuropsychiatric illness. On average, participants had a median of 1.0 year (SD=7.5 years) of formal music training. All were paid for their time and gave informed consent in compliance with a protocol approved by the Institutional Review Board at the University of Memphis.

### Speech stimulus continuum and behavioral task

We used a synthetic five-step vowel continuum previously used to investigate the neural correlates of CP (Oldfield, 1971) (Figure 1a). Each token of the continuum was separated by equidistant steps acoustically based on first formant frequency (F1) yet was perceived categorically from /u/ to /a/. Tokens were 100 ms, including 10 ms of rise/fall time to reduce spectral splatter in the stimuli. Each contained an identical voice fundamental (F0), second (F2), and third formant (F3) frequencies (F0: 150 Hz, F2: 1090 Hz, and F3: 2350 Hz). The F1 was parameterized over five equal steps between 430 and 730 Hz such that the resultant stimulus set spanned a perceptual phonetic continuum from /u/ to /a/ (Bidelman et al., 2013). Speech stimuli were delivered binaurally at 83 dB SPL through shielded insert earphones (ER-2; Etymotic Research) coupled to a TDT RP2 processor (Tucker Davis Technologies). Listeners heard 150-200 trials of each individual speech token during EEG recording. On each trial, they were asked to label the sound with a binary response (“u” or “a”) as quickly and accurately as possible (speeded classification task). Reaction times (RTs) were logged, calculated as the timing difference between stimulus onset and listeners’ behavioral response. Following their keypress, the inter-stimulus interval (ISI) was jittered randomly between 800 and 1000 ms (20 ms steps, uniform distribution) and the next trial was commenced.

**Figure 1:**
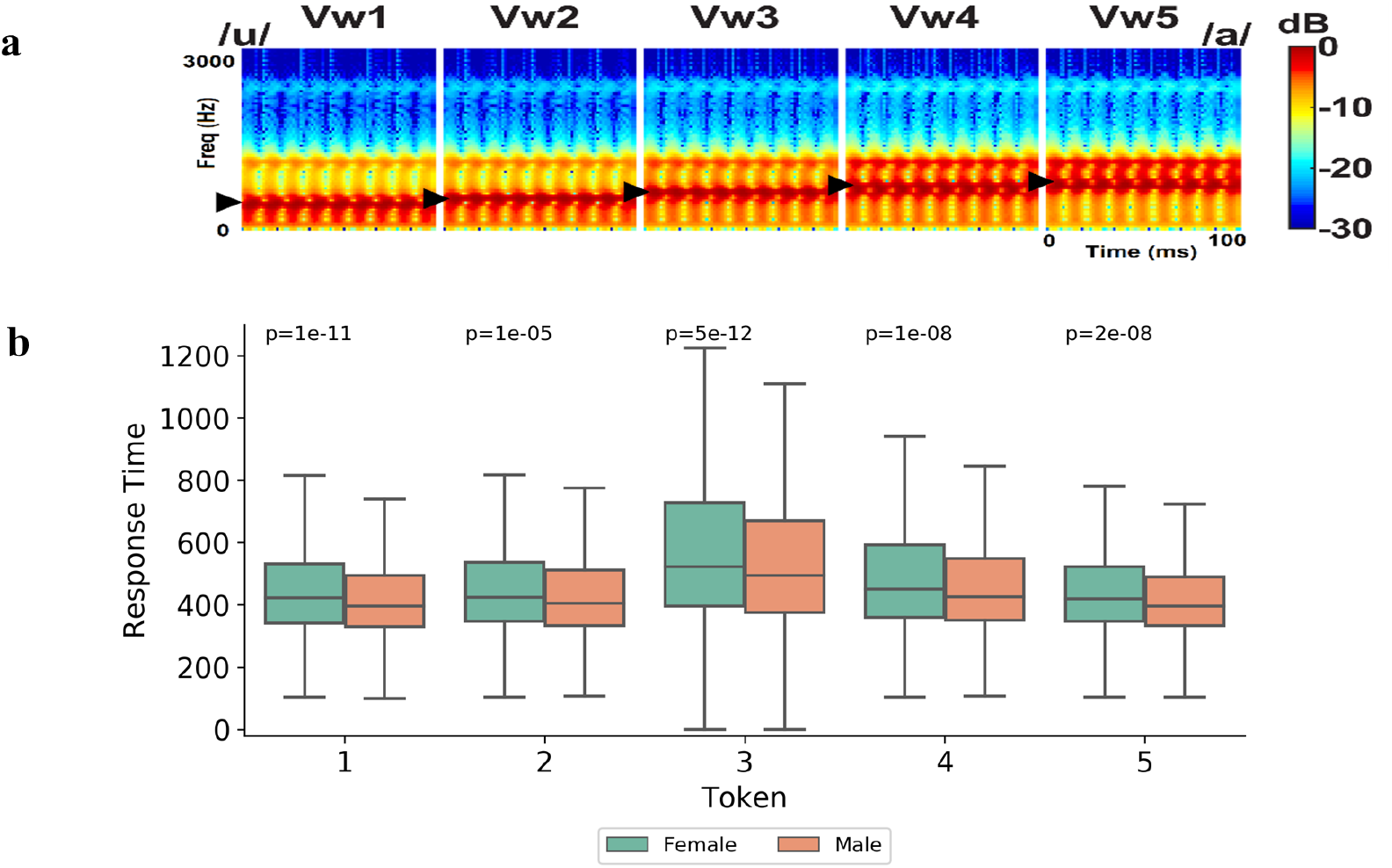
(a) Acoustic spectrograms of the speech stimuli: The stimulus continuum was created by parametrically changing vowel first formant frequency over five equal steps from 430 to 730 Hz, resulting in a perceptual-phonetic continuum from /u/ to /a/. (b) Token wise response times for auditory classification. Listeners are slower to label sounds near the categorical boundary (i.e., Token 3). Females had significantly slower RTs than males.

Our speech categorization task required listeners make a binary judgement on what they hear. As such, it is a subjective task that does not have true accuracy, *per se*. Consequently, we chose to decode RTs since they are a continuous, more objective measure that provides a much richer decoding of listeners’ behavior regarding the sound-to-label process.

### Behavioral data analysis

We adopted a Gaussian mixture model (GMM) with expectation-maximization (EM) to identify an optimal number of clusters/components (i.e., subgroups of listeners’ RTs) from the aggregate distribution of their RT speeds. Finding an optimal number of components with GMM is challenging. Here, we used brute-force and Bayesian Information Criterion (BIC) based approaches. In this exhaustive parameter search, the hyperparameters were: (1) Number of components (clusters), (ranges from 1 to 14), and (2) Type of covariance parameters. This process identified an optimal combination of four components with the unique covariance matrix. It was observed that 17% - 47% of the total trials in the speech identification task fell into three components. The 4th component had the fewest number of trials (1.6%). Based on the interpretation of RTs, we categorized these components as four clusters: Fast RT (120 - 476 ms), Medium RT (478 - 722 ms), Slow RT (724 -1430 ms), and Outliers (> 140 ms). Data from the outliers were discarded for further analysis given the low trial counts loading into this cluster. The boxplot in Figure 2 shows token wise RTs. Each speech token can be broken down into a combination of the three RT clusters, meaning that speech categorization speeds could be objectively clustered into fast, medium, slow (and outliers) responses via the GMM. These cluster divisions were then used in subsequent EEG analyses to determine if functional brain connectomics differentiated these perceptual subgroups.

**Figure 2:**
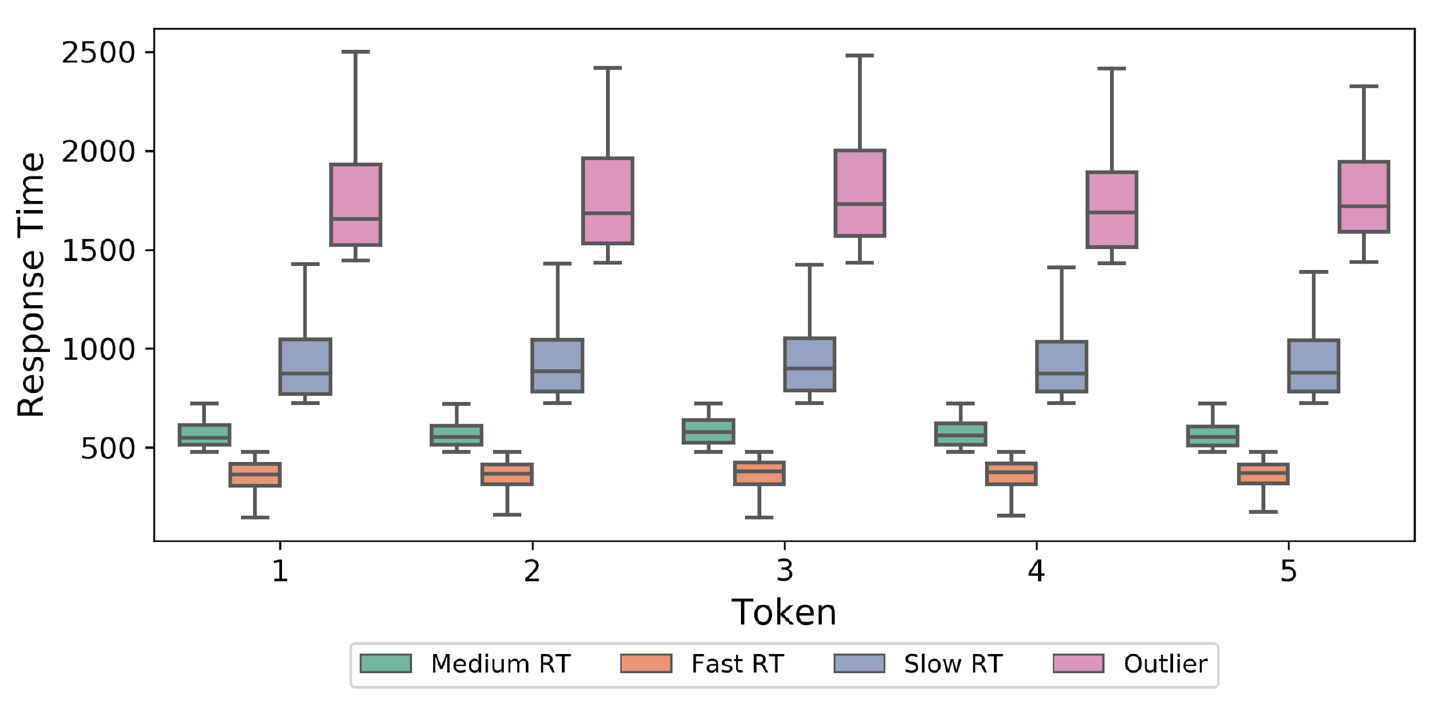
Token-wise RTs broken down by GMM component. Based on behavioral RTs, four clusters are evident that distinguish subgroups of listeners based on their speech identification speeds: Fast (120 - 476 ms), Medium (478 - 722 ms), Slow (724 - 1430 ms), and Outlier (> 1430 ms).

### EEG recording and preprocessing

EEG recording procedures were identical to our previous neuroimaging studies on speech categorization (Bidelman & Alain, 2015; Bidelman et al., 2013; Bidelman & Walker, 2017). Briefly, neuroelectric activity was recorded from 64 sintered Ag/AgCl electrodes at standard 10-10 locations around the scalp (Oostenveld & Praamstra, 2001). Continuous data were digitized using a sampling rate of 500 Hz (SynAmps RT amplifiers; Compumedics Neuroscan) and an online passband of DC-200 Hz. Electrodes placed on the outer canthi of the eyes and the superior and inferior orbit monitored ocular movements. Contact impedances were maintained < 10 kΩ during data collection. During acquisition, electrodes were referenced to an additional sensor placed 1 cm posterior to the Cz channel. Subsequent pre-processing was performed in BESA® Research (v7) (BESA, GmbH). Ocular artifacts (saccades and blinks) were first corrected in the continuous EEG using a principal component analysis (PCA) [48]. Cleaned EEGs were then filtered (bandpass: 1-100 Hz; notch filter: 60 Hz), epoched (−200-800 ms) into single trials, baselined to the pre-stimulus interval, and re-referenced to the common average of the scalp. This resulted in between 750 and 1000 single trials of EEG data per subject (i.e., 150-200 trials per speech token). Trials of same class were ensemble averaged per subject. This resulted in 105 data samples (=35 subjects x 3 RT classes) for further analysis. A schematic of the data pipeline is shown in Figure 3.

**Figure 3:**
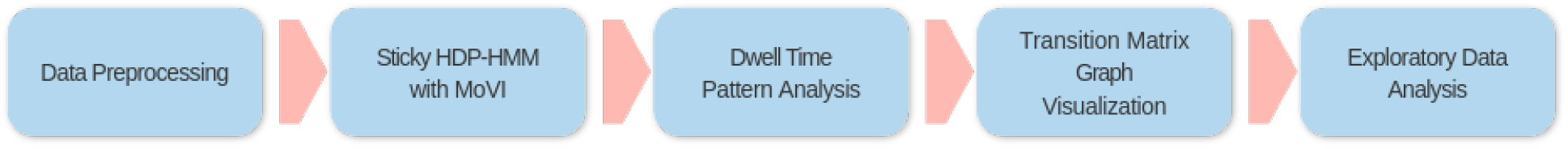
Schematic diagram of the processing pipeline. 64 channel EEG data was first preprocessed, and trialwise averaged. Sticky HPD-HMM with Memoized Variational inferance (MoVI) was then applied for data segenetation into functional brain microstates. Exploratory data analysis (e.g., dwell time pattern) was then used for further analysis and interpretation.

### Hierarchical Dirichlet Process Hidden Markov Models

We used the BNPY library (M. C. Hughes & Sudderth, 2014) in Python for spatiotemporal EEG data segmentation. This framework supports Bayesian nonparametric clustering and captures multidimensional, sequential, spatial, and hierarchical structures. To run inference on a dataset, BNPY requires an allocation model, a data-generation method, and the inference algorithm. The allocation model describes the generative process that allocates cluster assignments to individual data points. Here, we used HDP-HMM (Markov sequence models with an infinite number of states).

Observation models define a likelihood for producing data from cluster-specific density. We used Diagonal-Covariance Gaussian as observation models. However, the inference algorithm optimizes a variational-bound objective function. To achieve scalability, we focused on modern optimization-based approaches that can process batched data, particularly memoized variational inference. The mathematical definition and interpretation of Sticky HDP-HMM with Memoized Variational Inference (MoVI) are described in the appendix.

### Calculation of microstate and dwell time statistics

Dwell time quantifies the duration the EEG spends in a particular “functional microstate” before transitioning to another (Khanna et al., 2015; Miller et al., 2010). We quantified several widely used dwell time statistics including: (i) Duration: average duration a given microstate remains stable, (ii) Occurrence: the frequency of each microstate independent of its individual duration which reflects the activation trend of a potential neural source, (iii) Time coverage: fraction of total recording time for which a given microstate is dominant, (iv) Global variance: the variance explained by each microstate, (v) Transition probabilities: the transition probabilities of a given microstate to any other microstate (Khanna et al., 2015; Koenig et al., 1999).

## RESULTS AND DISCUSSION

### Micro-state-based analysis of the brain’s speech categorization

We adopted a data driven micro-state-based model for understanding the dynamics of speech categorization using HDP-HMM. The HDP-HMM starts with an infinite number of states. The Birth, Merge, and delete proposals are widely used to remove ineffective states [37]. To achieve an interpretable number of states, we varied the number of clusters from 5 to 30 (step 5) and observed the cluster probability. Figure 4 shows the number of clusters (K) vs. cluster probability. Each bar shows the number of data points that load in that specific cluster of a specific model (out of 5 models). The first cluster (0) contained most of the data points. First 10 clusters accounted for 93% of data points and cluster numbers above 10 had very fewer data points. Hence, we chose K=10 to perform our analysis.

**Figure 4:**
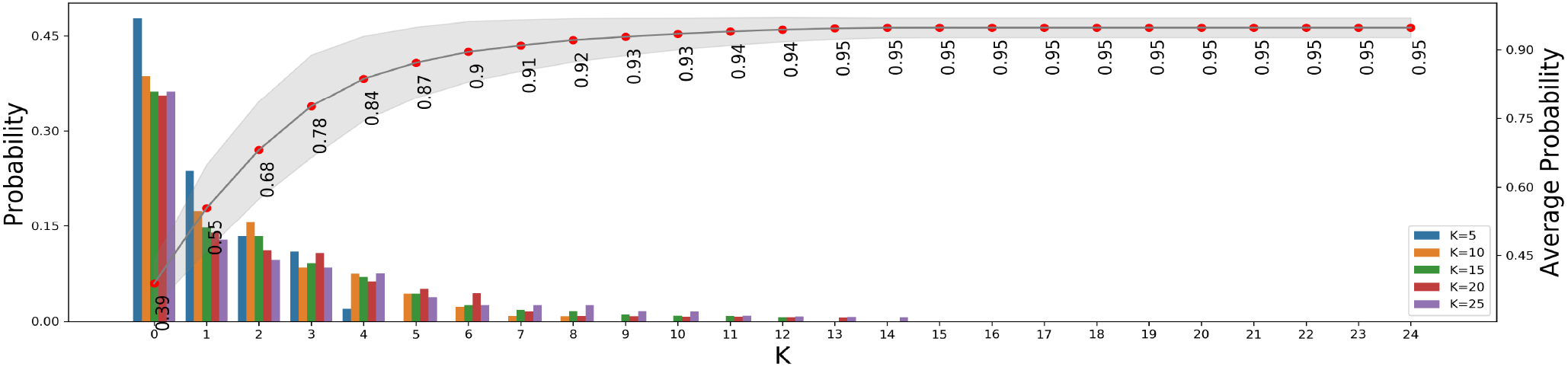
Probability of data points loading into each component.

### Computation and interpretation of micro-states underlying speech categorization

Using HDP-HMM with 10 states, we then segmented the EEG time series data. The time series data of a sample trial (Slow-RT of Subject #1) and dwell time pattern (tile and time-series) are shown in Figure 5. Each color of the tile represents the amount of time a particular state remains stable. The dwell-time shows the duration and pattern of transitions between microstates during the the 1 sec interval encompassing the sound presentation and subsequent categorization of a speech phoneme.

**Figure 5:**
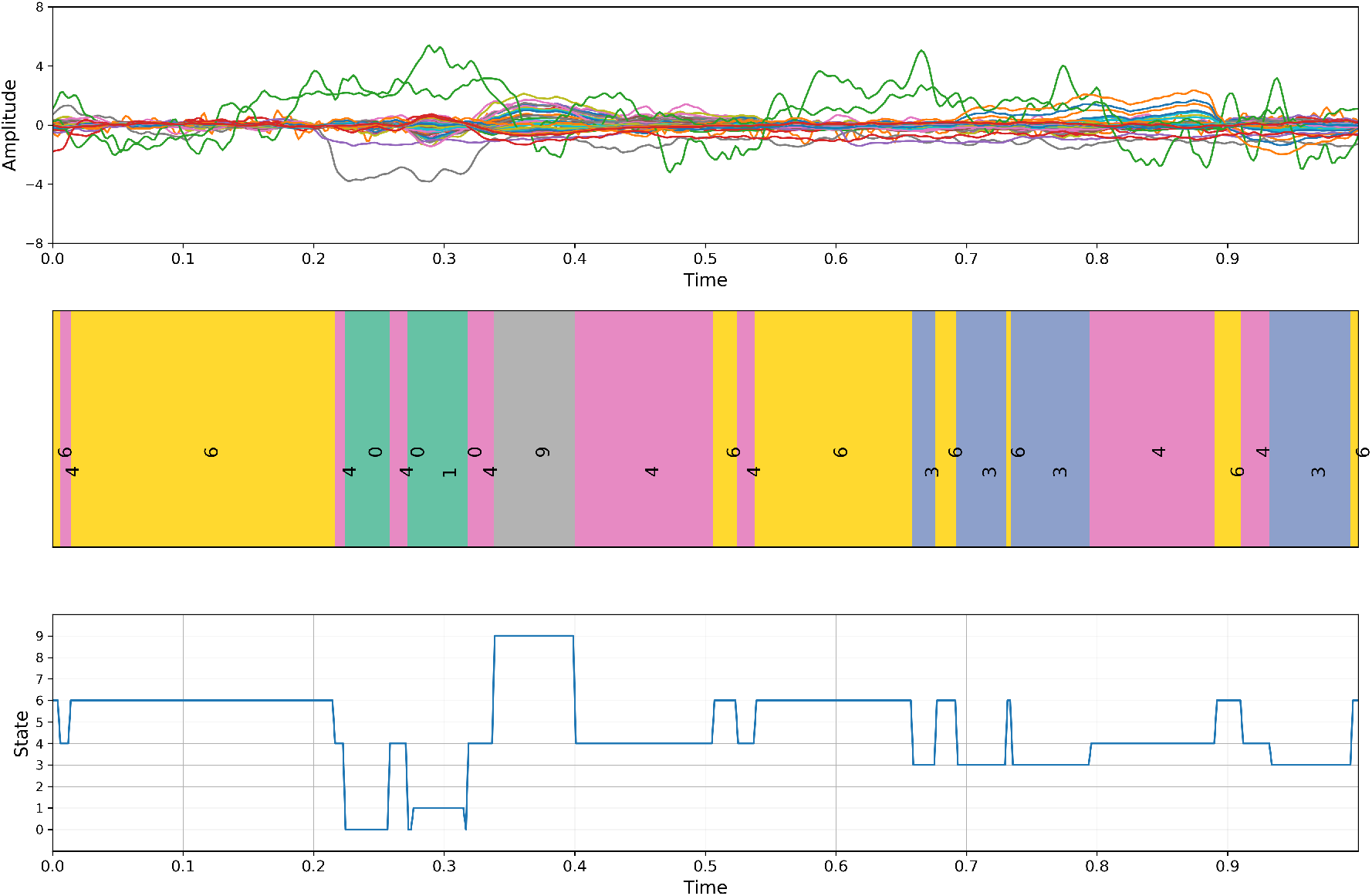
Top: Representative trial of 64 channel EEG data (Subject#1, slow RT). Middle: tile visualization of dwell pattern. Colors/numbers denote unique microstates. Bottom: time series visualization of dwell pattern illustrating the duration and transition between brain microstates.

The computed micro-states are interpretable based on what is known about the time course of the canonical evoked response to speech (i.e., event-related potential, ERP). For example, State 6 emerges and persists though the pre-stimulus (−200 - 0 ms) period, suggesting it represents a resting brain state prior to stimulus presentation. States 0 and 1 develop in the first 150 ms after stimulus presentation, co-occurring with the early exogenous waves (P1-N1) of the auditory cortical ERP which reflect stimulus encoding. Following, State 9 (a unique state) develops between 150 - 200 ms suggesting this segment reflects neural activity associated with the conventional P2 deflection, a wave associated with the formation of auditory perceptual objects and abstract categories. The return to State 6 between 300-500 ms suggests a return of the EEG to rest prior to the behavioral response. Finally, emergence of States 3-4, microstates which heavily loaded toward the end of the trial, suggests these states reflect processes related the preparation and/or execution of listeners’ motor response which logs their category decision. These data confirm that unique and highly interpretable functional states of brain activity (e.g., baseline resting, stimulus encoding, decoding, response selection) are readily decoded from the temporal dynamics of EEG during active speech perception.

We calculated trial-wise dwell time statistics to quantify different properties of microstates derived from HDP-HMM segmentation. Figure 6 shows the summary statistics of state-wise dwell time pattern. These analyses revealed different temporal extents in terms of the duration and frequency with which the EEG stayed in different states. Microstates 3, 4, and 6, for instance, occurred more often across the sample and persisted for longer durations. The duration of these states was also modulated by listeners’ RTs such that faster decisions were related to longer durations of microstates 3, 4, and 6. Interpreted in the context of the task, these results imply that faster categorization of speech at the behavioral level is associated with a longer and more dwell times within the putative resting state (microstate 6) and response selection (microstate 3 - 4). We found that slower RT decisions were associated with longer stays in microstate 0 compared to medium and fast RTs. Similarly, the early timing of microstates 0-1 lead us to infer they reflect stimulus coding (see Fig. 5). This suggests that brain activity spends a preponderance of time encoding stimulus features in trials having slower perceptual decisions. Collectively, these results can be summarized as three key:

**Figure 6:**
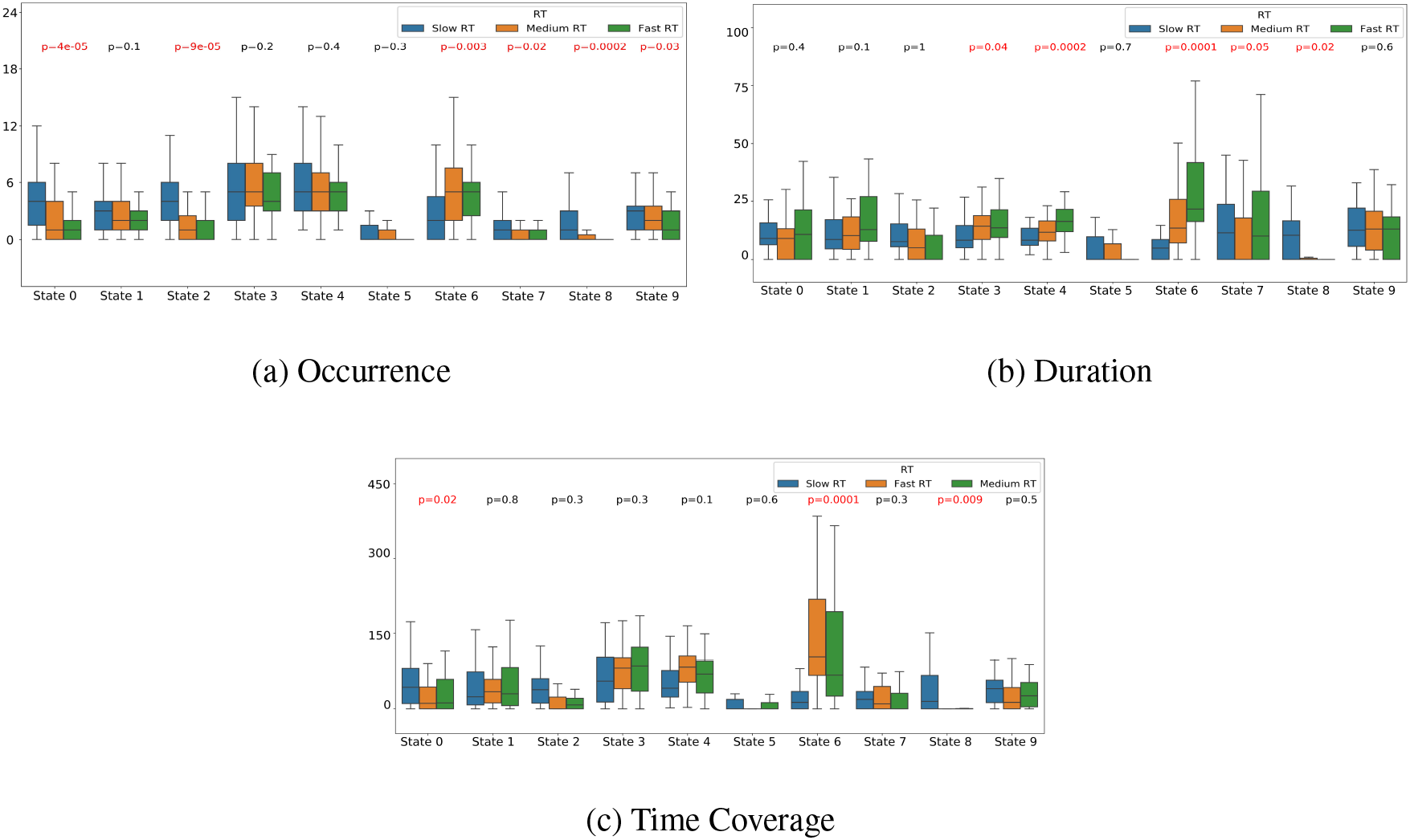
Bar plot visualization of dwell time statistics. Red values denote metrics that should significant (p < 0.05) differences between RT groups.

1. Trials which yield faster RTs are associated with the listener spending significantly less time in stimulus encoding (microstate 3), response selection (microstate 4), and resting state (microstate 6) patterns.. The brevity of time spend in both stimulus and response periods suggests the most rapid RTs in the behavioral task reflect fast guesses (i.e., “trigger” happy response).
2. RT wise, stimulus encoding, and response selection states occur with similar frequency overall.
3. Combining observation 1 and 2 suggests it is not important how frequently a listener stays in stimulus encoding and response selection state, *per se*. Rather, it is, how long they maintain those patterns brain states that is important to determining how fast they are able to label incoming speech sounds.
4. Trials with the faster RTs spend significantly more time in resting state, against suggesting an idling-like processing during the most rapid (likely guess) decisions.

### Relationship between speech RTs and transitional probabilities between brain microstates

Next, we examined the transition probabilities between different microstates in relation to the behavioral RTs. Figure 7 shows a heat map visualization of the transition matrix, which represents the likelihood that listeners’ changed from one state to another state. Diagonal elements represent the probabilities that a specific state remained stable (self-loop), whereas off-diagonal cells represent the likelihood of jumping from one state to another state. In general, states were highly stable and the transition probability between states was relatively low.

**Figure 7:**
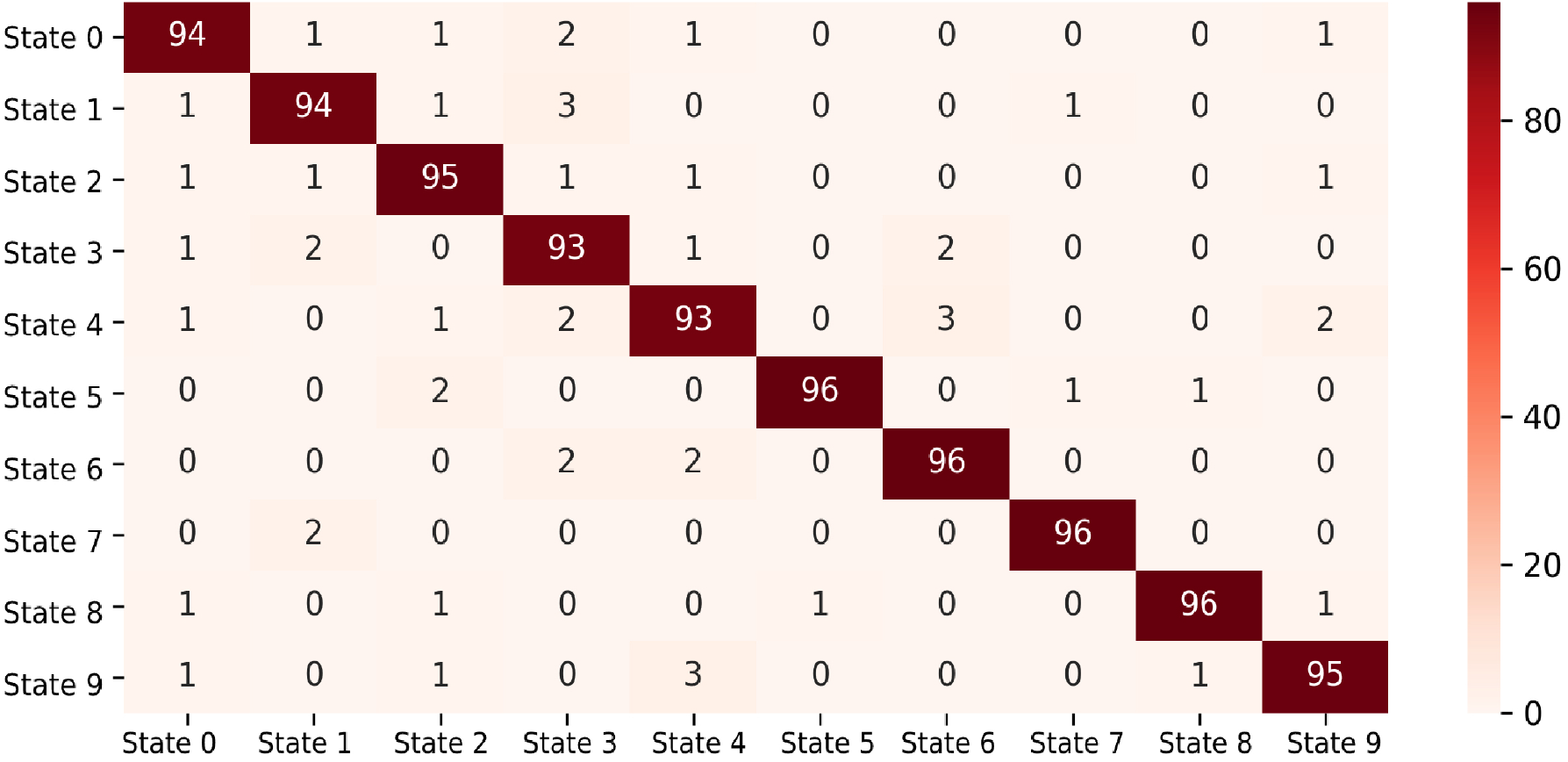
Transition matrix between brain microstates during the time course cateogrical speech processing. Probabilities are normalized to a range 0-100%.

We next asked whether the different behavioral groups (RT speeds) were associated with different frequencies of transitional probabilities. Figure 8 shows the frequency of state variations for the three RT groups computed from listeners’ transition matrices, along with the associated entropy for state variation. Results show a graded pattern from slow to fast RTs in both the frequency of state variation change and entropy. That this, when listeners were slow to label the speech stimuli, EEG entropy was high, suggesting more rapid progression through multiple functional states. In contrast, fast behavioral decisions were associated with lower entropy (i.e., less state variation), implying more stable functional states over time. These data corroborate the generally longer duration and time coverage of state dwell times for fast responders reported in Fig. 6.

**Figure 8:**
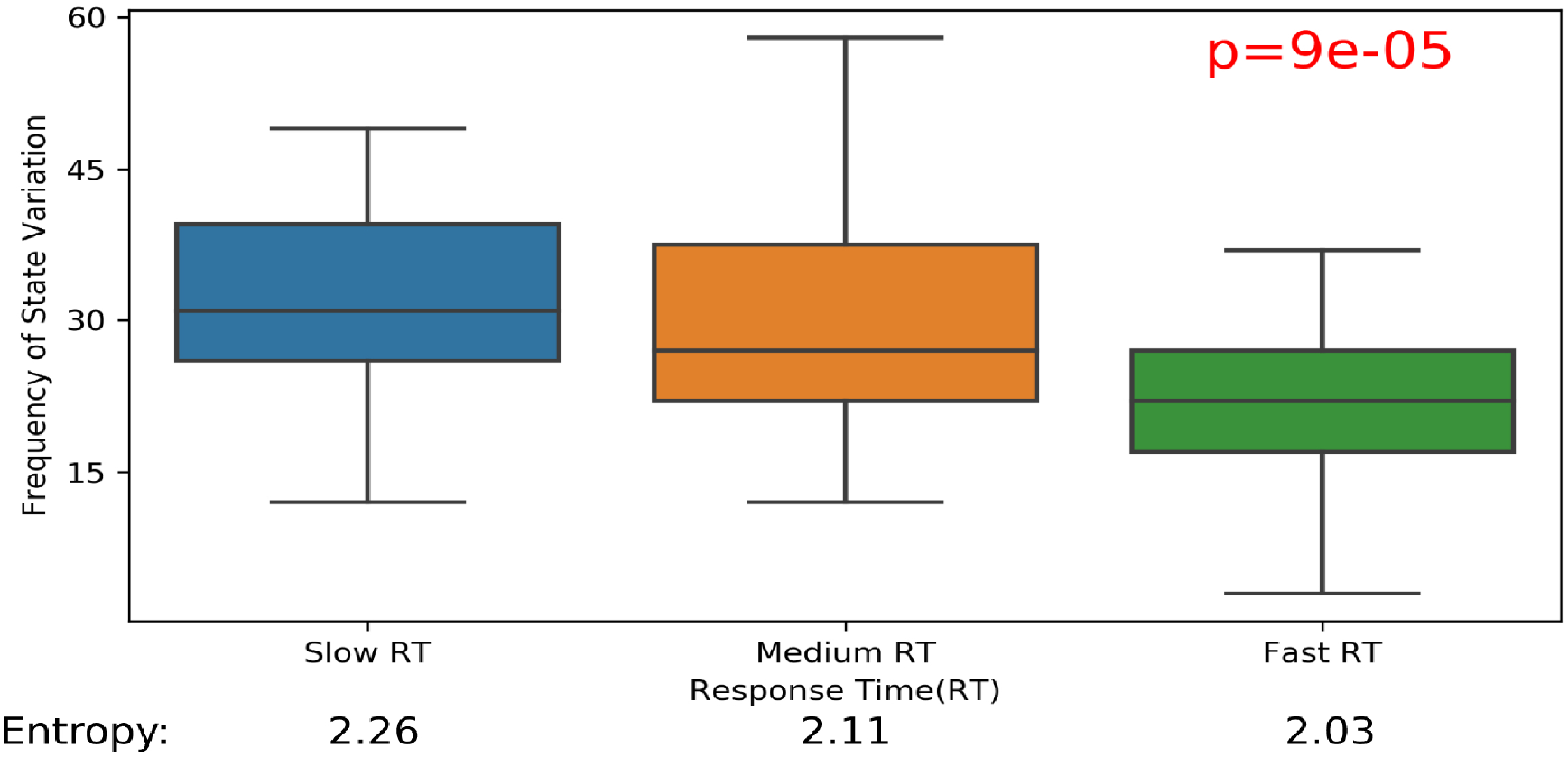
Frequency of EEG state change per RT group. State entropy is higher for slower compared to faster RTs suggesting poorer behavioral responses are associated with more rapid progression through multiple functional states.

Collectively, these findings suggest that perceptual speed in speech categorization is proportional to transition frequency. In states of slow speech perception decisions, the brain is more likely to change states rapidly, whereas faster decisions are accompanied by more stable patterns of brain activity. Moreover, entropy is higher for slower compared to faster RTs implying that the decision making process is more uncertain for slower behavioral responses, perhaps due to the more rapidly changing microstates we observe at the neural level.

### Differential networks underlying different RT groups

It is possible to compute the transition matrix from the Sticky HDP-HMM analysis from the transition matrix (see Figure 7). Figure 9 show a directed graph of the transition matrix for slow, medium, and fast responses in the speech CP task.

**Figure 9:**
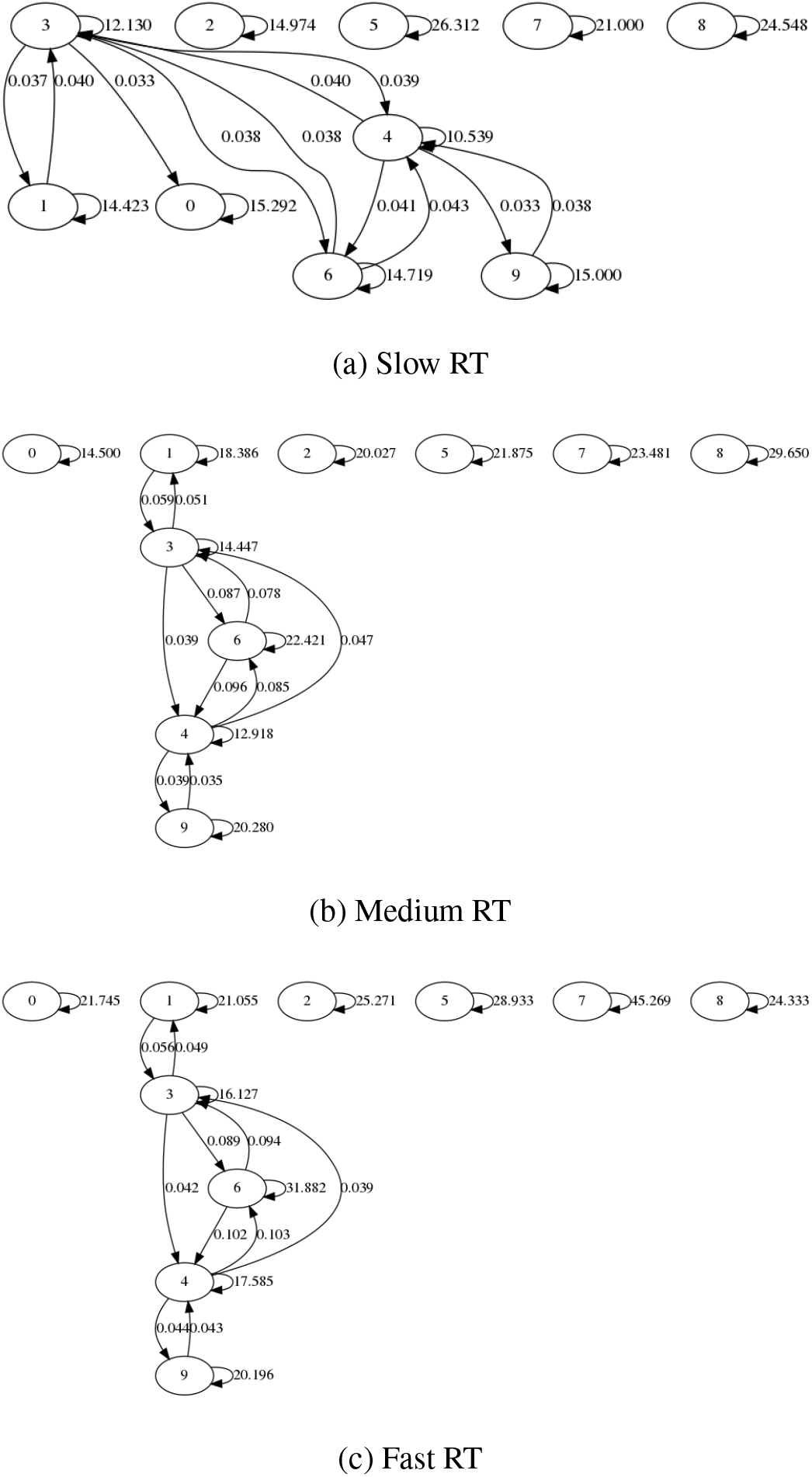
Graph visualization of transition matrix meta-analysis. Graphs relfect transition matrices (see Fig. 97) for differnet RTs in the speech categoriztion task. Each node represents one state. Self-loop nodes represent the average time a state remains stable in condition-specific (RTs) t rials. Edges represent state transition probabilities with direction inicated by arrows. Probabilities < 0.03 are discarded for better visualization with a smaller number of nodes.

These graph visualizations reveal the dynamic changes in brain states during rapid speech categorization and how different skill levels in the task (i.e., fast vs. slow responses) alter state dynamics. In general, we find that slower behavioral responses are associated with more pervasive exploration of different brain states. In contrast, faster perceptual RTs seem to be associated with a relatively compact oscillation among a more restricted set of states (e.g., microstate 1, 6, and 9). These findings support the notion that during rapid speech perception, slower perceivers utilize functional brain networks excessively (or inappropriately) whereas fast perceivers utilized the same neural pathways but with more restricted organization (Al-Fahad et al., 2020).

From the network structures presented in the Fig. 9, there appears to be a common delta (triangular) pattern in medium and fast RTs and slow RT seems to form a random graph. Graphs consisting of resting-state (microstate 6), stimulus encoding (microstate 3), response selection (microstate 4) which we argue represents a core decision-making default network (DN) underlying speech categorization. We observe that the primary cause of slower RTs during the fundamental process of sound identification is a longer time span in the DN. On the contrary, a more direct and rapid transition in the DN occurs during faster perceptual responses. In summary, our data suggest that the speed of speech categorization decisions is inversely proportional to the time listeners stay in the putative DN network.

### Intracranial sources underlying CP micro-states

Figure 10 shows topographic scalp map projections of the 10 microstates. We applied Classical Low Resolution Electromagnetic Tomography Analysis Recursively Applied (CLARA) [BESA Research (v7); BESA, GmbH] [51]–[53] to provide a qualitative description of the underlying brain sources that generate each state-specific scalp topography. CLARA renders more focal source reconstructions by iteratively reducing the source space during repeated estimations of the inverse solution. On each step, a spatially smoothed LORETA solution was recomputed, and voxels below a 1% max amplitude threshold are removed. This provided a spatial weighting term for each voxel of the LORETA image on the subsequent step. Two iterations were used with a voxel size of 7 mm in Talairach space and regularization (parameter accounting for noise) set at 0.01% singular value decomposition. CLARA activation maps were overlaid onto the BESA adult MRI template for visualization with respect to the brain anatomy. Figure 11 shows the CLARA source activations of what might be considered 4 primary states we have loosely identified as resting state (microstate 6), stimulus coding (microstate 1), category/auditory object formation (microstate 9), and response selection/motor planning (microstate 4).

**Figure 10:**
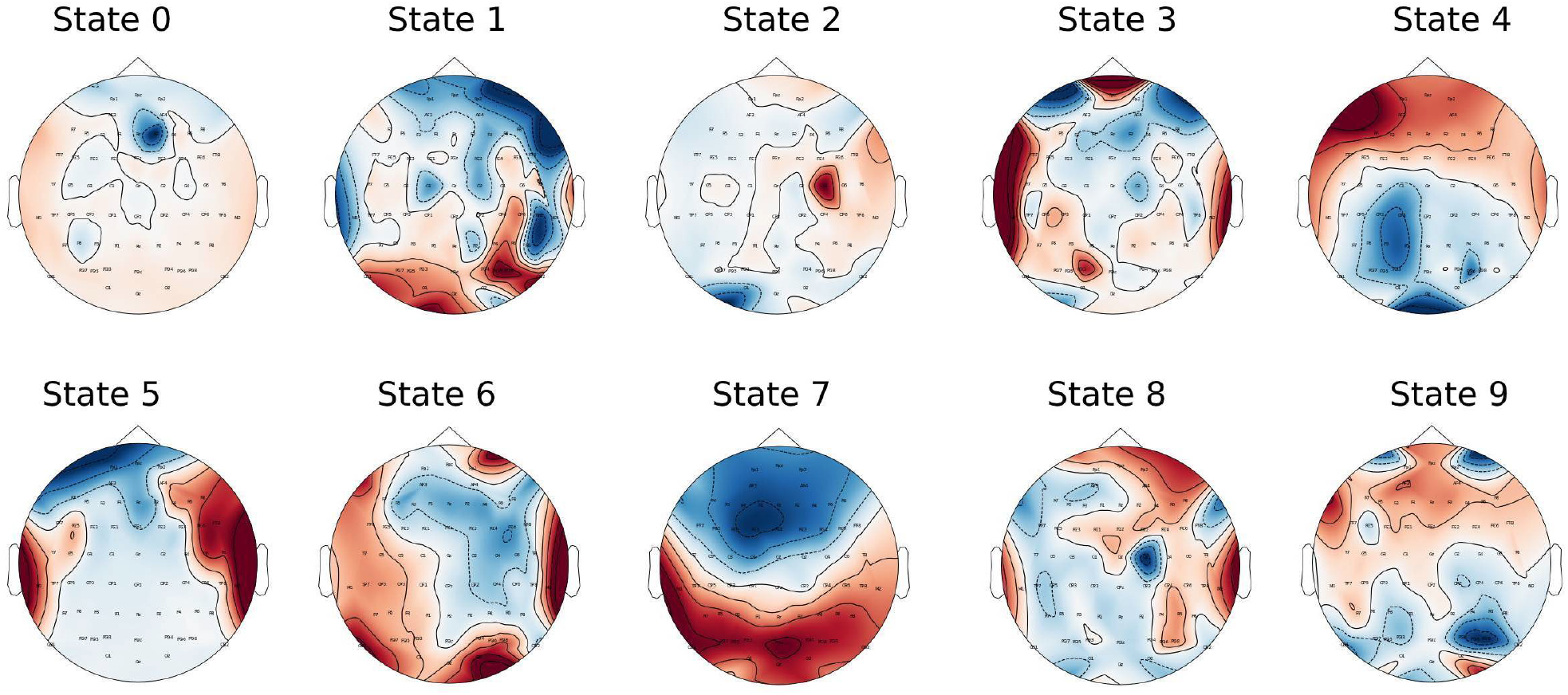
Topographic maps of the the 10 microstates underlying speech categorization.

These brain areas converge with both hypothesis and data-driven work which have shown a similar engagement of these regions in categorical decisions related to speech perception. For example, in our recent study (Mahmud, Yeasin, & Bidelman, 2021) applying machine learning techniques (e.g., neural classifiers, feature mining, stability selection) to full brain, source-reconstructed EEGs, we showed that 13 (out of 68) brain regions of the Desikan-Killiany (DK) (Desikan et al., 2006) atlas were dominant in describing listeners’ category labels for speech sounds during early periods of stimulus encoding (0–260 ms). Indeed, microstates 1 and 9 identified via Bayesian non-parametric analysis of response RTs, isolate these patterned activities to nearly identical brain areas (STG, IFG; see Fig.11) during the first 200 ms after the onset of the speech stimulus (see Fig. 5). Previous neuroimaging studies have shown that both STG (i.e., auditory cortex) and IFG (i.e., Broca’s area) are heavily involved in category decisions for speech (Bidelman & Walker, 2019). Similarly, we have shown that brain regions within the auditory-linguistic-motor loop including fronto-parietal, motor, and supramarginal (SMG) areas map to later decision stages of the categorization process within a time window of 300-800 ms (Bidelman, Pearson, & Harrison, 2021). Notable among our dynamic state analysis is the engagement of SMG during microstate 4 (Fig. 11) which occurred during a similarly late stage of the trial time course (300-700 ms, Fig. 5) putatively reflecting listeners’ response selection. SMG is involved in category decisions particularly those involving some ambiguity and may serve as a top-down modulator of speech processing that biases listeners’ decisions of heard speech (Bidelman et al., 2021).

**Figure 11:**
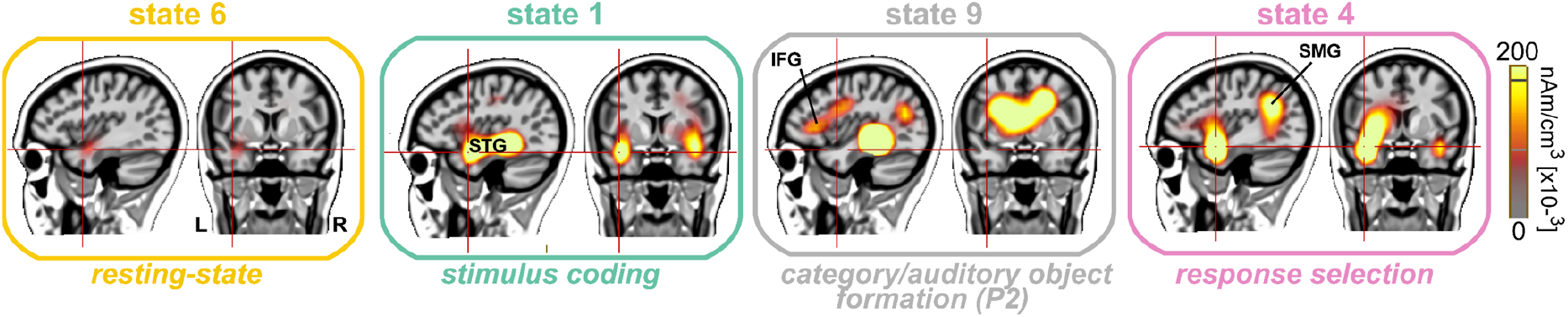
CLARA activation maps for selected brain microstates during categorial speech processing. STG = superior temporal gyrus (auditory cortex; stimulus coding). IFG = inferior frontal gyrus (home of important language regions like Broca’s area). SMG = supramarginal gyrus (implicated in lexical/semantic decisions including making phonological word choices).

We replicate and extend these prior findings by demonstrating a similar neural network describing listeners’ speed of categorical decision (i.e., RTs). It is worth noting that our microstate-based analysis here used an entirely different decoding approached applied to CP decision speeds (RTs) rather than listeners’ binary labels of speech sounds as in previous studies (Mahmud et al., 2021). Yet, the consistency of our findings across divergent studies, methods, and behavioral assays is striking. The similarity between regions and timing of their engagement across studies leads us to infer that the simple behavioral parameters of speech categorization—in terms both a listeners’ labeling accuracy and their speed are subserved by a relatively compact decision network (DN). Whether or not this neural circuit is entirely isomorphic in terms of processing both the accuracy and timing of the acoustic-phonetic mapping inherent to speech perception remains to be fully tested. However, in support of this proposition, recent neuroimaging studies have indeed shown that acoustic–phonetic conversion (cf. the label listeners place on speech sounds) and postperceptual decision (i.e., indexed by their RT) processes both localize to the same brain areas (Gow, Segawa, Ahlfors, & Lin, 2008). This raises the intriguing possibly that different aspects of behavior (at least with reard to perception) might be governed by the dynamics of a small, relatively compact brain network(s) rather than unique, domain-specific circuits, *per se*. This remains to be tested in future work. More broadly, certain developmental and neurocognitive disorders have been shown to impair the sound-to-meaning mapping process. Thus, tracking the temporal dynamics of functional brain microstates and how they might change over time with maturation or disease progression could provide new insight into receptive speech impairments in these clinical populations (Bidelman et al., 2017; Calcus Axelle et al., 2016; Werker & Tees, 1987).

## ACKNOWLEDGMENTS

This work was supported by the Department of Electrical and Computer Engineering, University of Memphis and the National Institute on Deafness and Other Communication Disorders of the National Institutes of Health under award number NIH/NIDCD R01DC016267 (G.M.B.).

## TECHNICAL TERMS

### Hierarchical Dirichlet Process Hidden Markov Models

Let EEG have n number of trials such that the data is represented as *x*_*n*_ = [*x*_*n*1_, *x*_*n*2_, *x*_*n*3_ . . . *x*_*nT*_]. Observation *x*_*nt*_ is a vector representing at time t and *x*_*nt*_ ∈ ℝ^*D*^. For 64 channel EEG data D = 64. The HDP-HMM explains this data by assigning each observation *x*_*nt*_ to a single hidden state *Z*_*nt*_. The chosen state comes from a countably infinite set of clusters K ∈ {1, 2, . . .}, generated via Markovian dynamics with initial state distributions *π*_0_ and transition distributions 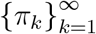.

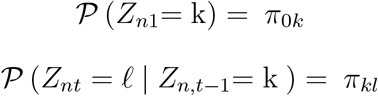

### Hierarchies of Dirichlet Processes

The number of states is unbounded under the HDP-HMM prior and posterior. The hierarchical Dirichlet process (HDP) shares states over time via a latent root probability vector *β* over the infinite set of states. The stick-breaking representation of the prior on *β* first draws independent variables *μ*_*k*_ = *Beta* (1, *γ*) for each state *k* and then set 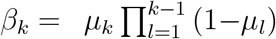. The *μ*_*k*_ can be interpreted as conditional probability of choosing *k* th state among states. The HDP-HMM generates transition distributions *μ*_*k*_ for each state *k* from a Dirichlet with mean equal to *β* and variance governed by concentration parameter *α*.

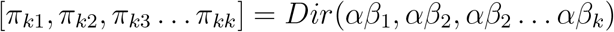

The *π*_*k*0_ is the starting probability vector with *π*_*k*0_∼ *Dir*(*α_o_β*_1_). Where *α_o_* ≫ *α*.

### Variational Inference

The inferential goal of HDP-HMM is to arrive at the posterior knowledge of top-level conditional probabilities.

*μ*_*k*_, HMM parameters: cluster probability *π*, cluster shape *ϕ* and cluster assignments *z* after observing data x. Parameter *μ*, *π* ,*ϕ* are considered as global parameter parameters because they generalize to new data sequences. The cluster assignments *z*_*n*_ is a local parameter specific to data sequence *x*_*n*_.

Variational methods frame posterior inference as an optimization problem [54]. Here, we seek a distribution q(*μ*, *π* ,*ϕ*) over the unobserved variables that is close to the true posterior i.e. q(*μ*, *π* ,*ϕ*, *z*) ∼ p (*μ*, *π* ,*ϕ*, *z* |*x*). We can re-present q(*μ*, *π* ,*ϕ*, *z*) as simpler factorized family q(*μ*, *π* ,*ϕ*, *z*) ≅ q(*μ*) q(*π*) q(*ϕ*) q(*z*). Inference algorithms update these parameters to minimize the Kullback-Leibler (KL) divergence. Equation for KL divergence is given by:

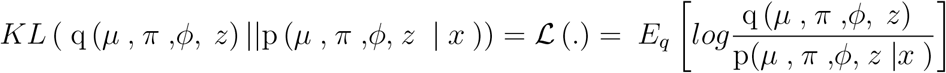

To get the best q(*) distribution, we need to optimize this objective function 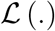. This equation can be simplified with four components.

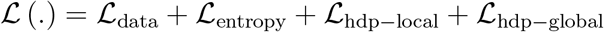

Here

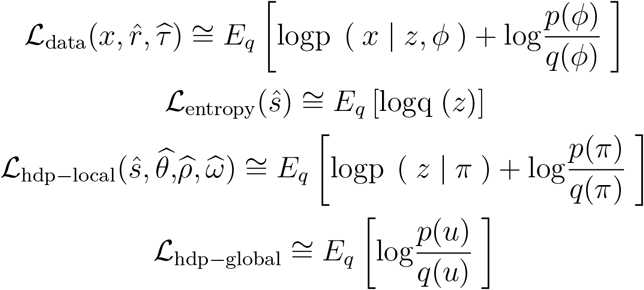

### Memoized and Stochastic Variational Inference

Common variational inference algorithms maximize 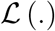 using coordinate ascent optimization. Here the optimal value of each parameter is kept fixed while optimizing other parameters. For the sticky HDP-HMM variational objective, each sequence is randomly assigned to one of *B* batches initially. The algorithm repeatedly and random visits batches one at a time. Each full pass through the complete set of *B* batches a is considered as a lap. At each visit to batch *b*, sticky HDP-HMM perform a local step for all sequences *n* in batch *b* and then a global step. The batch optimization of 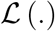 is possible by exploiting the additivity of statistics *M, S*. For each statistic, this algorithm track batch-specific quantity *M ^b^*, and a summary of whole-dataset 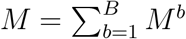.

After a local step at batch *b*, yields 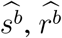 and update 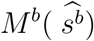 and 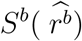, increment each whole-dataset statistic by adding the new batch summary and subtracting the summary stored in memory from the previous visit and store (or memoize) the new statistics for future iterations. It is possible to evaluate 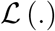 at any point during memoized execution except 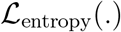 term. To compute it, a (K + 1) × K dimensional matrix *H^b^* is tracked at each batch *b*. Where:

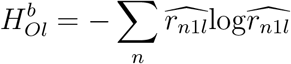

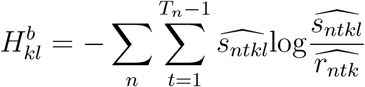

For the whole dataset, the entropy matrix is defined as:

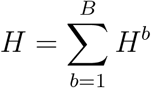

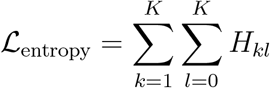

